# Progesterone receptor activation regulates sensory sensitivity

**DOI:** 10.1101/2023.08.04.552037

**Authors:** Suchitra Joshi, John Williamson, Shayan Moosa, Jaideep Kapur

**Author notes:** Correspondence Suchitra Joshi MSc, PhD, Department of Neurology, University of Virginia, Health Sciences Center, PO Box: 801330, Charlottesville, VA 22908.

## Abstract

Women develop chronic pain during their reproductive years more often than men, and estrogen and progesterone regulate this susceptibility. We tested whether brain progesterone receptor (PR) signaling regulates pain susceptibility. During the estrous cycle, animals were more sensitive to pain during the estrus stage than in the diestrus stage, suggesting a role for reproductive hormones, estrogen, and progesterone. We measured the pain threshold daily for four days in ovariectomized, estrogen-primed animals treated with progesterone. The pain threshold was lower 2 days later and stayed that way for the duration of the testing. A specific progesterone-receptor (PR) agonist, segesterone, promoted pain, and mice lacking PR in the brain (PRKO) did not experience lowered pain threshold when treated with progesterone or segesterone. PR activation increased the cold sensitivity but did not affect the heat sensitivity and had a small effect on light sensitivity. Finally, we evaluated whether PR activation altered experimental migraine. Segesterone and nitroglycerin (NTG) when administered sequentially, reduced pain threshold but not separately. These studies have uncovered a pain-regulating function of PRs. Targeting PRs may provide a novel therapeutic avenue to treat chronic pain in women.

## Introduction

Female reproductive hormones regulate pain susceptibility rendering them more susceptible to chronic pain conditions, including migraine, tension-type headache, neuropathic pain, and temporomandibular pain (Silberstein, 2000; Yunus et al., 2000; Torrance et al., 2006; de Mos et al., 2007; Vincent and Tracey, 2008; Fillingim et al., 2009; Finnerup et al., 2021) and have a lower threshold for experimental pain than men (Fillingim et al., 2009; Bartley and Fillingim, 2013). Increased pain susceptibility in women often emerges following puberty, which marks cyclic hormonal fluctuations. Furthermore, the pain symptoms frequently worsen during the perimenstrual phase when hormone levels are low (Houghton et al., 2002; Sherman and LeResche, 2006; Martin, 2009; Tan et al., 2020) and improve during pregnancy when hormone levels are high (LeResche et al., 2005; MacGregor, 2007).

Progesterone exerts acute analgesic actions. Its administration a few hours before testing increased the pain threshold in animals with neuropathic pain or inflammatory allodynia (Verdi et al., 2013; Coronel et al., 2014; Hornung et al., 2020). Progesterone metabolite allopregnanolone likely exerts these analgesic actions by potentiating GABAergic inhibition (Coronel et al., 2016a), since agents that potentiate GABAergic inhibition are potent analgesics (reviewed in (Munro et al., 2013). However, the pain-suppressing actions of progesterone are not uniform. It did not alleviate pain induced by sciatic nerve cuffing (Dableh and Henry, 2011) and was also ineffective in suppressing perimenstrual migraine headaches in a small study (Somerville, 1971; Anne MacGregor, 2004). Progesterone also has other neuronal targets, including progesterone receptors (PRs) (Mani and Oyola, 2012; Kapur and Joshi, 2021). PR activation increases neuronal excitability, whereas allopregnanolone lowers it (Kapur and Joshi, 2021). Also, unlike the rapid actions of allopregnanolone, PR activation induces gene expression with effects emerging in several hours to days (Kapur and Joshi, 2021). This complexity could explain the discrepant results of progesterone treatment in treating chronic pain conditions. However, unlike allopregnanolone, the pain regulation by PRs remains poorly understood.

PRs are ligand-activated transcription factors, and their regulated effects emerge slowly but last longer (Brinton et al., 2008; Mani and Oyola, 2012). We have described PR activation’s slow, delayed excitatory effect (Joshi et al., 2018; Shiono et al., 2021). Since the pain paroxysms are thought to occur due to increased firing of the neurons in the pain pathway or because of a lowering of their firing threshold, the excitatory action of PR activation may be critical for the pain pathophysiology.

We performed this study to evaluate the role of PRs in regulating sensory sensitivity. We found that PR activation increased sensitivity to mechanical and thermal stimuli, and primed animals to the pain-promoting effects of nitroglycerin (NTG).

## Materials and methods

### Materials

Segesterone, a progesterone receptor agonist sold under the brand name Nestorone, progesterone, and allopregnanolone were purchased from Sigma-Aldrich. Nitroglycerin was obtained from American Reagent. All the other common chemicals were also obtained from Sigma-Aldrich.

### Animals

Animals were handled according to a protocol approved by the University of Virginia Animal Care and Use Committee (ACUC), compliant with the ARRIVE guidelines. All the animals had ad libitum access to food and water and were maintained on a 12-hour light and 12-hour dark cycle (lights on at 6 AM, lights off at 6 PM). Four to five animals were housed together. The experiments were performed with adult female (50-70 day-old) C57Bl6 mice or female mice lacking progesterone receptor expression. Mice with a floxed first exon of *Pgr* (PR^fl/fl^) were a kind gift from Dr. M. Luisa Iruela-Arispe (University of California, Los Angeles, CA) (Hashimoto-Partyka et al., 2006; Joshi et al., 2018). We and others have characterized these mice before (Goddard et al., 2014; Stephens et al., 2015; Joshi et al., 2018). These mice were crossed with nestin-Cre mice (B6.Cg-Tg(Nes-Cre)1Kln/J, The Jackson Laboratory # 003771) to generate animals lacking PR expression in the neurons and glia (brain-specific PR deletion, PRKO). The colony was maintained by breeding the PR^fl/fl^-cre^+ve^ male mice with PR^fl/fl^-Cre^-ve^ females. We will refer to PR^fl/fl^-cre^+ve^ mice as PRKO and PR^fl/fl^-Cre^-ve^ as wild-type (WT) mice throughout the manuscript. All the testing was performed between 9 AM to 2 PM.

### Estrous cycle monitoring

Vaginal smears of adult female C57Bl6 mice were obtained daily between 8 AM to 10 AM. The cytology of the wet smears was checked under a light microscope to determine the percentage of nucleated cells, non-nucleated corinified cells, or small round leukocytes. The animals in estrus phase (smear contained primarily cornified cells) and diestrus phase (presence of mostly leukocytes in the smear) were used for behavioral testing and blood collection for hormone measurements.

The serum progesterone levels (n=4) were measured using an ELISA assay (# K025-H1/H5 Arbor Assays, Ann Arbor, MI; detection range 50 to 3200 pg/mL). The intra-assay variation was 7% in these studies. The estradiol levels (n=4) were also measured using an ELISA assay (#ES180S-100 Calbiotech, detection range 3 to 300 pg/mL). The intra-assay variation was 5.9% in these assays.

### Ovariectomy and hormone treatment

Bilateral ovaries were removed under isofluorane anesthesia (Shiono et al., 2021). The experiments were performed after 10-12 days of recovery.

The animals were primed with 17β-estradiol (10 μg/animal, subcutaneous), and 2 days later with progesterone (10 mg/kg, subcutaneous, sc, n=6). The control animals received vehicle, 20% β-hydroxycyclodextrin (n=6). This progesterone treatment caused acute increase in serum progesterone levels (40-80 ng/mL, n=4), but the levels were back to baseline (3-5 ng/mL, n=4) 16 hr after the injection. Another cohort of C57Bl6 females were treated with PR agonist segesterone (10 mg/kg, sc, n=10, Sigma) or vehicle (20% β-hydroxycyclodextrin, n=9). A third cohort of C57Bl6 mice were treated with allopregnanolone (THP, 10 mg/kg, sc) or vehicle (DMSO), n=9 each. PRKO females and littermate wild-type animals were also treated with vehicle or segesterone (n=7 each) or progesterone (n=7 each). The animals were randomly assigned to the treatment groups.

### Evaluation of mechanical pain threshold

Mechanical pain sensitivity was tested using manual Von Frey monofilaments (Deuis et al., 2017). Briefly, the animals were individually placed in Plexiglas boxes (5” X 5” X 7”) with a wire mesh floor. The testing was performed after 30 min of acclimatization. The midplantar surface of each of the hind paws was tested thrice with von Frey filaments presented in an ascending order (0.4, 0.6, 1.0, 2.0, 4.0, 6.0, 8.0, 10.0 g). The filament was applied perpendicularly to the paw until the filament buckled slightly and held for 3 sec. Each stimulation is separated from the previous by 10 sec to prevent sensitization. A quick withdrawal of the paw and/or its licking marked a positive response. The filament that produced a positive response for 3 out of the presentations was considered as the threshold (Fig. 1B).

**Figure 1:**
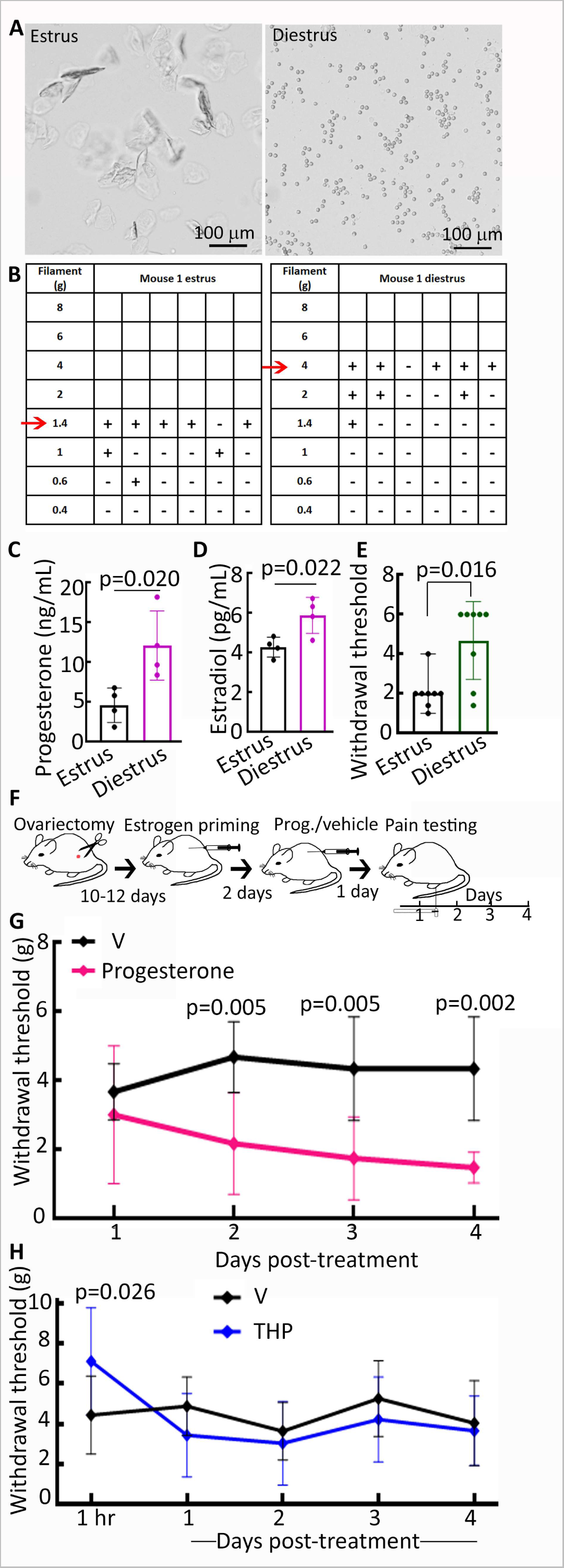
Mechanical pain sensitivity during estrous cycle and following progesterone treatment. **(A)** Representative images showing vaginal cytology of estrus and diestrus stages, the estrus stage was characterized by the abundance of cornified cells whereas the leukocytes were plenty during diestrus stage. **(B)** Responses to the presentation of von Frey filaments in ascending order in a representative animal tested in estrus and diestrus phases; + represents a positive response and – represents no response. The red arrow marks the mechanical pain threshold. **(C and D)** Mean and SD of the circulating progesterone and estradiol levels respectively in animals in estrus and diestrus stages (n=4), The p-values show results of the student’s t-test. **(E)** The pain threshold in animals in estrus and diestrus stages. The values represent mean ± SD of the Von Frey filament weight that produced a response on both the hind paws at least 50% of the times, n= 8, p-value represents the results of Wilcoxin matched-pairs signed-rank test. The same set of animals was tested during the two phases. **(F)** A schematic showing treatment and testing schedule. Adult C57Bl6 female mice were ovariectomized and primed with estrogen (17β-estradiol 10 μg, sc). Two days later the animals received progesterone (10 mg/kg, sc) or vehicle (20% β-hydroxycyclodextrin). The same group of animals was tested daily. **(G)** Mean and SD of the mechanical pain threshold in progesterone- or vehicle-treated mice, n=6 each, the p-values of Holm-Šídák’s multiple comparisons test are shown on the graph. **(H)** Effect of allopregnanolone (THP) on mechanical pain threshold, n=9 each, the p-value of Sidak’s multiple comparisons showing significant differences at 1 hr in the vehicle and THP-treated mice is shown.

### Evaluation of thermosensitivity

Cold sensitivity was measured by the latency to escape from a cold floor (15°C). The animals are kept in a custom-built Plexiglas chamber (5 X 5 X 7 inches) with a peltier-controlled cooling plate forming half the floor of the chamber. The remaining half of the floor was elevated (2.5 cm) to form an escape platform. The animals were acclimatized to the chamber for 5 min on the day before the testing. On the day of the experiment, the animals were introduced to the chamber on the cooling plate side and the peltier turned on to cool the plate from room temperature (23-25°C) to 15°C over a period of 3 sec. The time to jump away from the cold floor to the elevated platform was measured. To prevent injury to the animal, any animal that did not jump away in 30 sec was manually removed and the latency marked as 30 sec. Each animal was tested twice with 5 min interval during which the animal was returned to the home cage. Mean latency to climb onto the elevated platform was determined for each animal on each testing day (n=7).

Warm sensitivity was determined using the tail flick test (Schildhaus et al., 2014). The animals were restrained in a cylindrical tube and 2 min later, 3 cm of the end of the tail was dipped in hot water maintained at 48±1°C. The latency to flick the tail was recorded, and as done for the cold sensitivity assay, the assay ended at 30 sec. Each animal was tested twice with a 5 min interval between tests, and average latency was determined (n=7).

### Evaluation of light sensitivity

The light sensitivity was evaluated using a light-dark box (n=10), with a light (15 X 11 X 20 cm) compartment with transparent walls and floor, connected to a dark chamber of the same dimensions but with black walls through a 4 X 4 cm opening in the wall. The animals were introduced to the light compartment and allowed to explore both the compartments for 7 min. The movement of animals was video recorded and analyzed post-hoc using AnyMaze software. The time in light and dark compartments and the latency to the first entry to the dark compartment were evaluated.

### Nitroglycerin treatment

To induced migraine-like central sensitization, NTG 5, 10, or 15 mg/kg (n=7 each dose) or vehicle (n=7) were intraperitoneally injected. A 5 mg/ml stock of NTG (in 30% alcohol, 30% propylene glycol, and water, American Reagent) was diluted 1:1 with saline and then injected to animals, 2x the body weight for 5 mg/kg dose and 6x the body weight for 15 mg/kg dose. The control animals received injections of the vehicle diluted 1:1 with saline. The animals were randomly assigned to the four groups. Paw withdrawal threshold using Von Frey filaments was evaluated to assess the effect of NTG.

### Statistical analysis

Graphpad Prism 9 was used to perform statistical comparisons. The data are plotted as mean ± standard deviation. Wilcoxon matched-paired sign rank test was used to evaluate differences in pain threshold during estrous cycle. Student’s t-test and ordinary one-way ANOVA with post-hoc Šídák’s multiple comparisons test or Dunnet’s multiple comparison test were used to evaluate whether the treated and untreated animals differed from each other. The differences were considered significant when the p-value was less than 0.05.

## Results

### Pain susceptibility fluctuates during the estrus cycle

Women experience elevated pain sensitivity during the perimenstrual period. We assessed sensitivity to mechanical pain using manual Von Frey monofilaments (Fig. 1B) during two estrous cycle phases determined by vaginal cytology (Fig. 1A). The progesterone and estrogen levels were higher in the diestrus than in the estrus phase (Fig. 1C, 1D, n=4, t(6)=3.079, p=0.02) for progesterone and *t*(6)=3.081, p=0.022 for estrogen, student’s t-test). We tested five animals in the estrus stage first and subsequently in the diestrus stage, and three animals in reverse order. Animals in the estrus phase responded to a smaller force than those in the diestrus phase (Fig. 1B, 1E, n=8, p=0.016, Wilcoxin matched-pairs signed-rank test). Thus, similar to the heightened perimenstrual pain susceptibility in women, mice were also susceptible to mechanical pain during the low hormone level estrous phase. lowered pain threshold during the estrus stage could be due to the slow progesterone effect observed on its withdrawal or estrogen withdrawal. We focused on the PR-mediated slow impact of progesteronene.

### Lasting effects of progesterone lowered pain susceptibility

The animals were ovariectomized and primed with 17β-estradiol to avoid the confounding effects of endogenous hormonal fluctuations (Fig. 1F). The animals received a single injection of progesterone or vehicle two days after estradiol treatment and subsequently tested on four days (Fig. 1F, n=6 each). A day after progesterone administration, the paw withdrawal threshold was similar in the progesterone- and vehicle-treated mice (Fig. 1G). However, two days after the progesterone treatment, the animals’ pain threshold was lower than pretreatment, and the difference persisted (F(7, 40)=5.449, p=0.0002, ordinary one-way ANOVA, please see p values for the Holm-Šídák’s multiple comparisons test in the graph). On the other hand, the threshold remained stable for four days of testing in the vehicle-treated mice.

Thus, progesterone exerted a slow-emerging pain-promoting effect, which could be due to allopregnanolone withdrawal or PR activation. Allopregnanolone is a progesterone metabolite, and because of rapid metabolism, its levels closely follow those of progesterone (Tuveri et al., 2008; Kimball et al., 2020). Thus allopregnanolone withdrawal could lower the pain threshold on days 2-4 following progesterone treatment, alternately progesterone could activate PR -regulated cellular signaling with slow-onset but lasting effects. We allopregnanolone’s role from that of PR-signaling in regulating pain susceptibility.

### Allopregnanolone withdrawal and pain susceptibility

Ovariectomized and estrogen-primed animals were treated with allopregnanolone (10 mg/kg) or vehicle; the acute effects were evaluated an hour after its administration and the lasting effects were evaluated for four subsequent days (n=9 each). We observed a treatment effect compared to the vehicle (F(9, 80)=3.160, p=0.0026, ordinary one-way ANOVA). Šídák’s multiple comparisons tests showed that the allopregnanolone elevated pain threshold 1 hr after THP administration (Fig. 1H, p=0.026), but the two groups were indistinct on days 1-4 of testing (Fig. 1H). Thus, allopregnanolone withdrawal did not explain the delayed pain-promoting effect of progesterone.

### PRs and regulation of pain susceptibility

We found that PR-mediated effects emerge 24-48 hrs after progesterone administration, consistent with the genomic action of these receptors (Joshi et al., 2018; Shiono et al., 2021). If PRs regulated the pain-promoting effect of progesterone, their deletion should block its effects, and their activation should reduce the pain threshold. We first evaluated the effect of progesterone treatment in the mice lacking PR expression in the nervous system (PRKO, PR^fl/fl^-cre+ve) and used littermate wild-type (WT, PR^fl/fl^-Cre-ve) mice as controls. The pre-progesterone pain threshold in the WT and PRKO mice was similar (*t*(12)=0.000, p>0.99, student’s t-test), indicating that when the endogenous hormonal fluctuations are eliminated, the two groups were comparable.

Progesterone did not change the pain threshold in the PRKO mice (n=7, F(4, 30)=0.5975, p=0.67, ordinary one-way ANOVA). In contrast, progesterone reduced the pain threshold in the WT mice (Fig. 2A, n= 7, F(4, 30)=3.380, p=0.021 ordinary one-way ANOVA). Post-hoc comparisons revealed a substantial reduction in the pain threshold on days 3 and 4 compared to the pretreatment pain threshold (p=0.011 and p=0.017, respectively, Dunnett’s multiple comparisons test).

**Figure 2:**
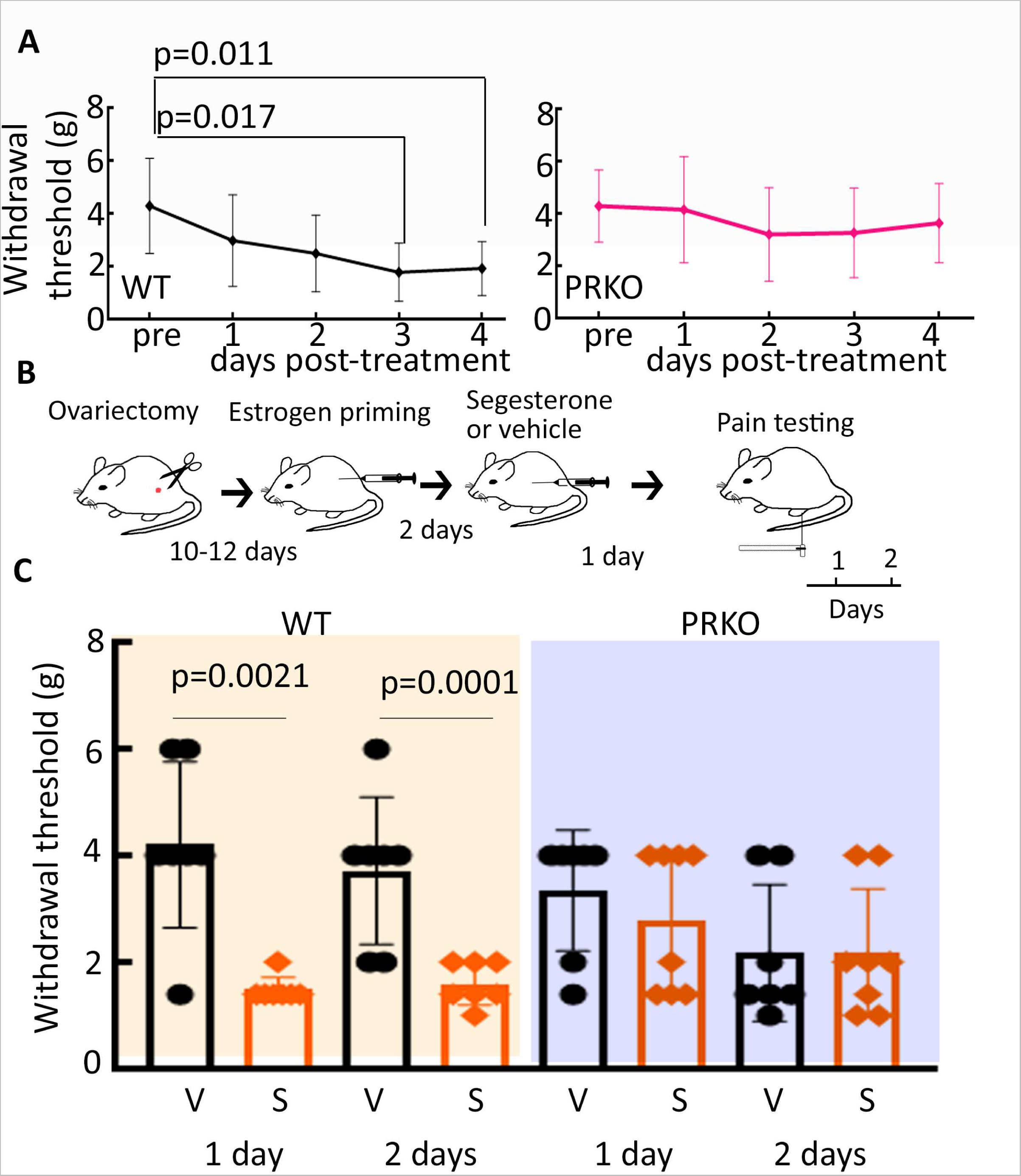
PR activation and mechanical pain sensitivity. **(A)** Mechanical pain threshold in the PRKO and littermate WT mice treated with progesterone, n= 7 each, the p-values of post-hoc Dunn’s multiple comparisons are shown. **(B)** A schematic showing segesterone or vehicle treatment of the animals. **(C)** The mechanical pain threshold in the WT and PRKO mice treated with vehicle or segesterone, n=7 each for WT, 7 vehicle-treated PRKO, and 8 segesterone-treated PRKO, the p-values of the post-hoc Šídák’s multiple comparisons test are shown.

We used segesterone, a specific PR agonist to confirm further the role of PRs in regulating pain sensitivity. Segesterone was injected in the ovariectomized, and estradiol-primed WT and PRKO mice with (10 mg/kg, sc, n=7 WT and 8 PRKO). Control animals received vehicle (20% β-hydroxycyclodextin, n=7 WT and 7 PRKO). Both groups were tested (Fig. 2B). The segesterone-treated control mice withdrew their paws from lower force filaments than the vehicle-treated mice (Fig. 2C). The main treatment effect was observed. Posthoc comparison showed lowering of pain threshold was seen on both the days of the testing (F(3, 24)=12.27, p<0.0001, ordinary one-way ANOVA, please see the post-hoc Šídák’s multiple comparisons test results in the graph). In contrast, the paw withdrawal threshold was similar between the vehicle- and progesterone-treated PRKO mice (Fig. 2C, F(3, 26)=1.481, p=0.2427, ordinary one-way ANOAV). Taken together, these studies revealed a pain-promoting effect of PR activation.

We treated intact cycling C57Bl6 mice with segesterone because ovariectomy has short and long-term effects, which could confound the PR activation effects described above (McElroy and Wade, 1987). Animals in the estrus stage were injected and tested for two days (n=7 each). The pain threshold in the segesterone-treated intact, cycling animals was lower than those that received vehicle injections (day 1, V: 5.343 ± 1.739 and S: 1.314 ± 0.3625 and day 2, V: 4.343 ± 2.797 and S: 2.200 ± 1.296, n=7 each, F(3, 24)=7.679, p=0.0009 ordinary one-way ANOVA, post-hoc Šídák’s multiple comparisons test results p=0.0006 1 day V vs S, and p=0.07 2 days V vs S).

### PR activation and thermal pain sensitivity

PR activation lowered heat but not cold sensitivity in female mice. We compared the sensitivity to warmer (48°C) and colder (15°C) temperatures in the segesterone and vehicle-treated mice to evaluate whether PR activation also regulated thermal nociception. We measured the latency to escape the cold floor. The vehicle and segesterone-treated mice were distinct in their sensitivity to the cold floor (Fig. 3A, n= 7 each, F(3, 24)=4.964, p=0.0081, ordinary one-way ANOVA). Post-hoc comparisons revealed that the escape latency was substantially shorter on the 2^nd^ day in the segesterone-treated animals than those treated with a vehicle (p=0.006, Šídák’s multiple comparisons test).

**Figure 3:**
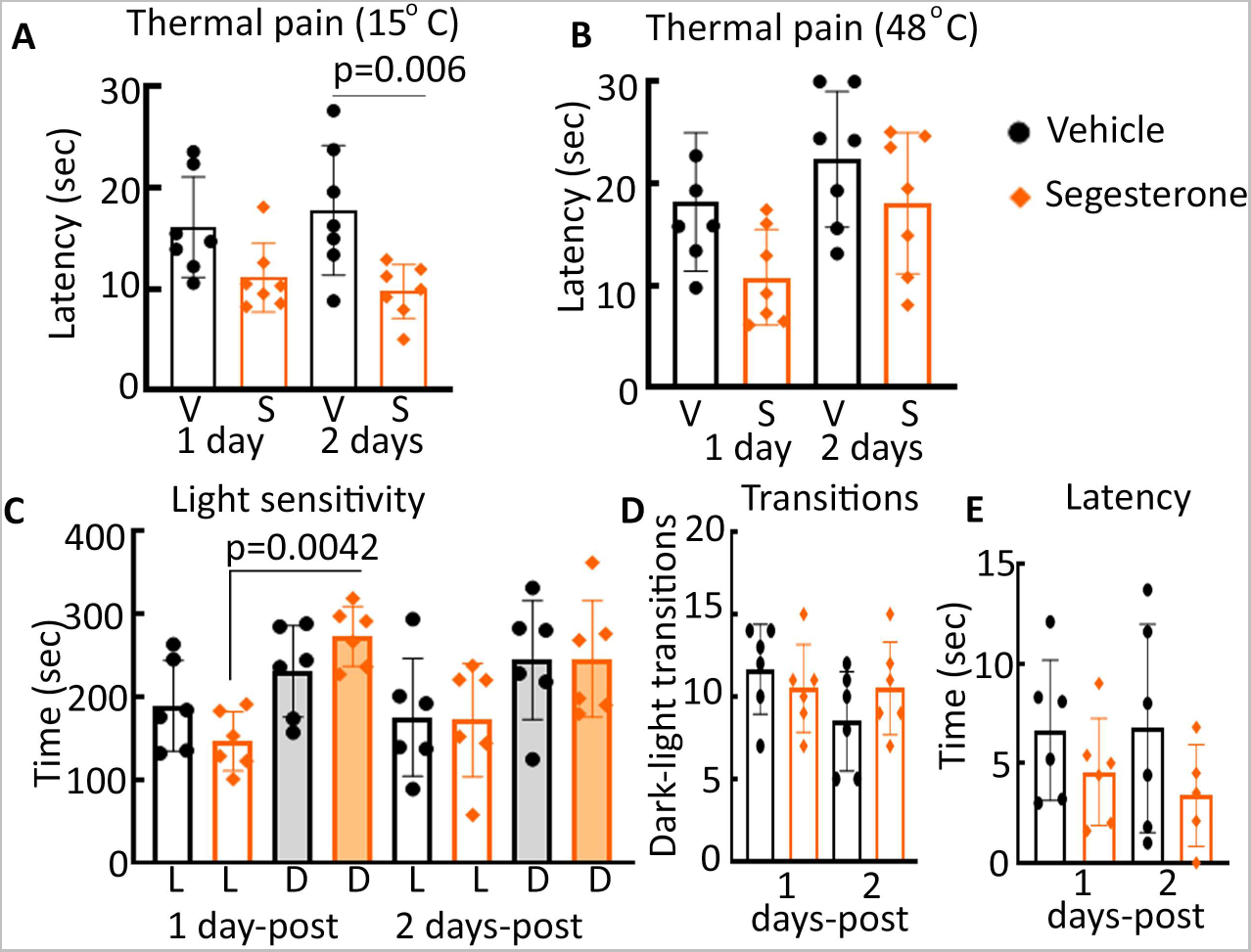
The effect of PR activation on thermal pain and light sensitivity. **(A)** Latency to escape (mean ± SD) the cold floor in mice treated with vehicle or segesterone, n=7 each, p-value of the post-hoc Šídák’s multiple comparisons test are shown. **(B)** The tail-flick latency in the vehicle- and segesterone -treated mice, n=7 each. **(C)** The time spent in the light (L) and dark (D) chambers respectively after treatment with segesterone (10 mg/kg)or vehicle, n=6 each. **(D)** The number of transitions between the light and dark compartments. **(E)** Latency to the first entry into the dark chamber.

We used a tail flick assay to measure warmer temperature sensitivity following segesterone treatment (Fig. 3B, n=7 each). Although an overall effect of the treatment was observed (F(3, 24)=4.060, p=0.0182, ordinary one-way ANOAVA), post-hoc comparisons did not uncover significant differences between the vehicle- and segesterone-treated animals (Fig. 3B, p=0.0757 and p=0.3840 respectively, Šídák’s multiple comparisons test). Thus, PR activation increased sensitivity to cold but did not alter the sensitivity to warmth.

### PR activation and light sensitivity

Migraines and tension type headaches are often associated with photophobia (Charles, 2018). Therefore, to extend the findings that uncovered a role of PRs in regulating tactile and thermal sensitivity, we also evaluated if PR agonist segesterone altered light sensitivity. The animals’ behavior in the light-dark box was tested after administration of segesterone (n=6) or vehicle (n=6). An increase in light sensitivity following segesterone treatment could lead to increased time in the dark compartment, fewer transitions between the two compartments, and/or a shorter latency to enter the dark compartment. Thus, we compared these parameters in the vehicle- and segesterone-treated animals. An overall effect of the treatment was observed on the time spent in the dark and light compartments (Fig. 3C, F(7, 40) = 3.365, p=0.0065, ordinary one-way ANOVA). Post-hoc comparisons revealed that segesterone-treated mice spent more time in the dark compartment than that in the light compartment a day after the treatment (p=0.0042, post-hoc Sidak’s multiple comparison test) whereas the vehicle-treated mice spent similar times in the two compartments (p=0.8002, post-hoc Sidak’s multiple comparison test). The time spent in the dark and light compartments did not differ in either of the groups on the 2^nd^ day. The dark-light transitions did not differ between the vehicle and segesterone-treated mice (Fig. 3D; F(3, 20)= 1.315, p= 0.2974 ordinary one-way ANOVA) and the latency to enter the dark compartment also did not differ between the two groups (Fig. 3E; F(3, 19)= 1.092, p= 0.3767, ordinary one-way ANOVA). Thus, PR activation had only a mild effect on light sensitivity.

### Gender- and age-specific effects of PR activation on mechanical pain threshold

Gender differences that disproportionately affect females exist for various pain conditions. To evaluate whether PR activation regulated pain in males, we treated adult male mice with segesterone and evaluated the paw withdrawal threshold. The threshold seemed comparable between segesterone- and vehicle-treated males (Fig. 4A, n=7). An overall effect of the treatment was observed (F(3, 24)=3.333, p=0.036, ordinary one-way ANOVA), but post-hoc comparisons did not reveal differences between the two treatment cohorts on either of the days (at 1 day p=0.1106 and at 2 days p=0.1106 Šídák’s multiple comparisons test). Thus, PR activation did not lower the mechanical pain threshold in adult males.

**Figure 4:**
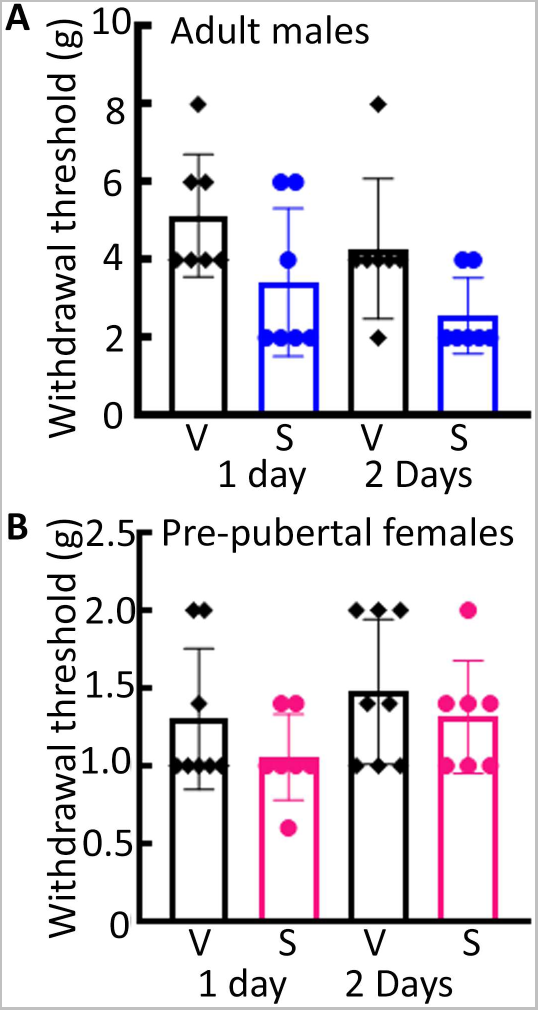
The effect of PR activation on pain susceptibility in males and juvenile females. **(A)** The mechanical pain threshold in adult male mice treated with vehicle or Nestorone, n=7 each, F(3, 24)=3.333, p=0.036 ordinary one-way ANOVA. Post-hoc Dunn’s multiple comparison test did not reveal significant differences between vehicle and Nestorone -treated mice (1 day-post: p=0.1106 and 2 days-post p=0.1106 Šídák’s multiple comparisons test). **(C)** The pain threshold in juvenile, pre-pubertal female mice (23 day-old), n=8 vehicle and 7 Nestorone -treated mice (F(3, 26)=1.357, p= 0.2777, ordinary one-way ANOVA).

Puberty marks the onset of cyclic hormonal changes in females, accompanied by the emergence of gender differences in the prevalence of clinical pain. We evaluated whether the onset of puberty increased the sensitivity to the effects of PR activation. The vaginal opening occurred in 30-32 day-old C57Bl6 females. Hence, we evaluated the effect of segesterone in 23-26 day-old females, before the onset of puberty (n=8 vehicle and 7 segesterone). Overall, the mechanical pain threshold was lower in the juvenile animals compared to that in the gonadally-intact adult females (*t*(15)=7.095, p<0.0001, student’s t-test). However, the pain threshold did not differ between the vehicle- and segesterone-treated mice, and it also remained stable over the 2 days of testing (Fig. 4B, F(3, 26)=1.357, p= 0.2777, ordinary one-way ANOVA). Together these studies showed that PR signaling was not associated with pain regulation in adult males and juvenile females.

### Nitroglycerin (NTG) reduced the mechanical pain threshold

The regulation of pain sensitivity by PR activation raised the possibility that PR signaling may also regulate pathological pain. Therefore, in the subsequent studies, we evaluated whether PRs regulated migraine-accompanying sensory hypersensitivity. NTG has been extensively used to induce migraine-like alterations in experimental animals and it reduces mechanical pain threshold (Tassorelli et al., 2003; Bates et al., 2010; Pradhan et al., 2014; Casili et al., 2020; Caudle et al., 2020). Ovariectomized and estrogen-primed mice were treated with three doses of NTG (5, 10, 15 mg/kg, n=7 each) and the pain threshold was evaluated one and four hours later. An overall effect of the treatments was observed (Fig. 5A, F(7, 48)=4.631, p=0.0005 ordinary one-way ANOVA). Post-hoc comparisons revealed that 10 mg/kg and 15 mg/kg doses of NTG reduced the pain threshold at both the time points compared to the threshold in vehicle-treated mice at the respective time points (at 1 hr: p=0.004 vehicle vs 10 mg/kg NTG, p=0.005 vehicle vs 15 mg/kg NTG; at 4 hr: p=0.007 vehicle vs 10 mg/kg NTG and p=0.025 vehicle vs 15 mg/kg NTG, Šídák’s multiple comparisons test).

**Figure 5:**
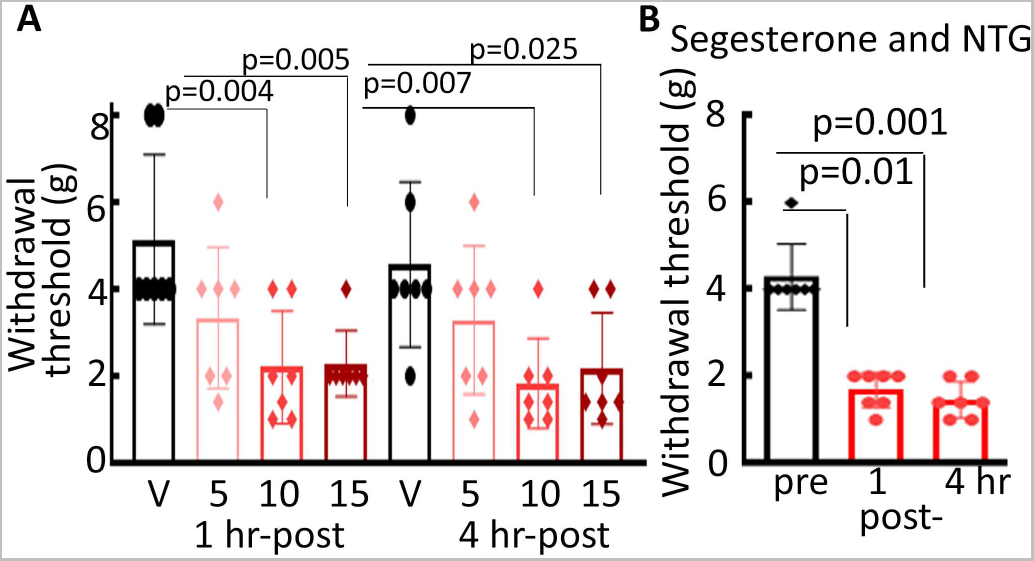
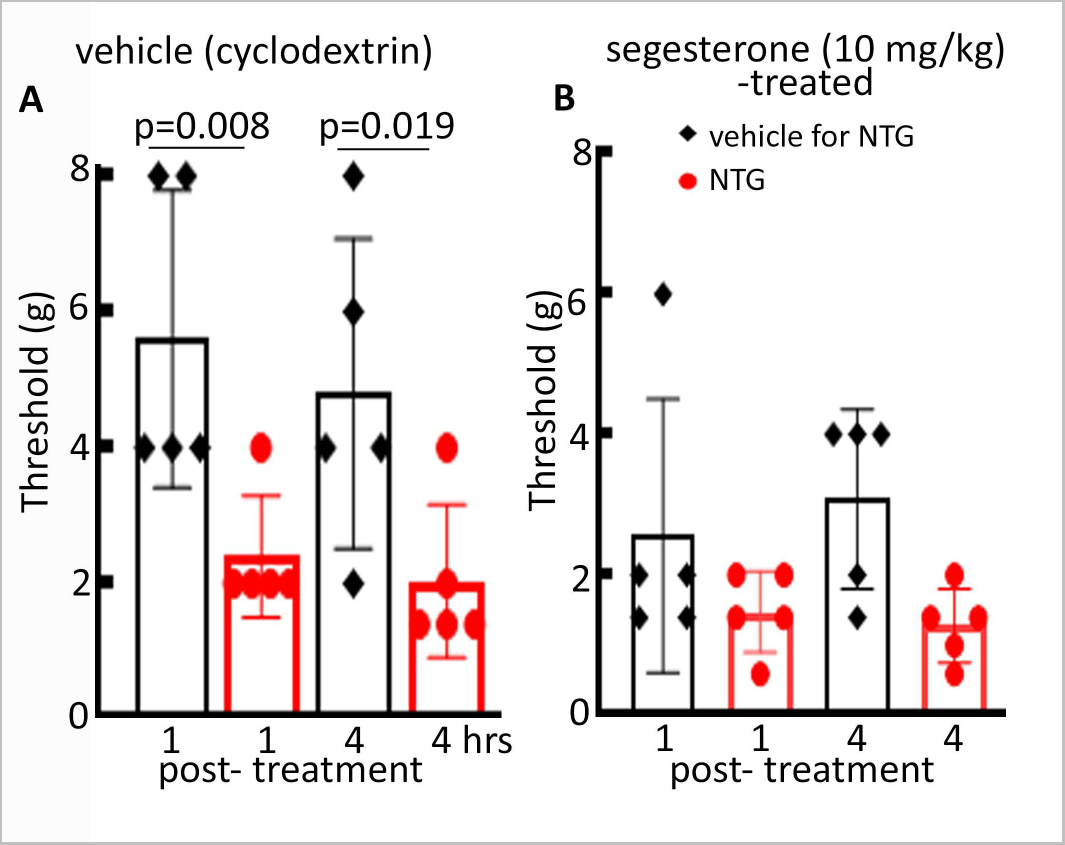
PR activation primed the animals for nitroglycerin-induced reduction in mechanical pain threshold. **(A)** The mechanical pain threshold in mice treated with NTG at 5 mg/kg, 10 mg/kg, or 15 mg/kg or with vehicle. The testing was done at 1 hr and 4 hr after the injections, n=7 each, the p-values of the post-hoc Sidak’s multiple comparisons are shown. **(B)** The mechanical pain threshold in animals that received low dose segesterone (3 mg/kg) followed by low dose NTG (5 mg/kg). The threshold was determined 2 hours prior to, and 1 and 4 hours after NTG administration, n=7, p<0.0001 ANOVA, the p-values of the post-hoc Šídák’s multiple comparisons test are shown.

Subsequently, we evaluated the effect of segesterone pretreatment on the NTG-induced pain. The animals were treated with segesterone (10 mg/kg) and a day later received NTG. However, the pain threshold in the segesterone+NTG-treated mice (n=6) was similar to that in segesterone+vehicle-treated mice (n=5) (supplementary figure 1, F(3, 18)=1.363, p=0.2860 ordinary one-way ANOVA).

### Synergistic action of segesterone and NTG

Since segesterone treatment itself reduced the pain threshold, it may not have caused a further reduction in the pain threshold following NTG administration. However, if segesterone and NTG exert a synergistic action, lower doses of these agents, at concentrations that individually don’t alter the pain threshold, could lower the pain threshold when administered together. To test, this we first evaluated the effect of 3 mg/kg segesterone on the mechanical pain threshold. This dose did not impact the pain threshold (4.29 ± 0.76, n=7), which was similar to that in animals treated with vehicle (3.83 ±1.9, n=7, *t*(12)=0.5956, p= 0.5625, student’s t-test).

We then evaluated the effect of low dose NTG (5 mg/kg) administration on the animals treated with low dose (3 mg/kg) segesterone. NTG was administered one day after segesterone. NTG reduced the pain threshold in these animals (Fig. 5B, n=7, F(2, 18)=56.82, p <0.0001, ordinary one-way ANOVA). The post-hoc comparison revealed that the pain threshold 1 and 4 hr after NTG administration was lower than that measured before its injection (p<0.0001 at both the time points, Šídák’s multiple comparisons test). Thus, a combination of low dose segesterone and NTG exerted a synergistic action to reduce the pain threshold.

## Discussion

We have uncovered a pain-promoting effect of PR activation in female mice. Progesterone and PR receptor agonist segesterone reduced pain threshold and this effect was blocked by brain-wide deletion of PRs. Furthermore, PR activation also primed the animals to painful effects of nitroglycerin.

Pain susceptibility or its perception varies across menstrual cycle in women (Hellström and Anderberg, 2003; Martin, 2009); it is higher during late luteal/early follicular phases, when the hormone levels are low compared to that during the follicular or ovulatory phases when the hormone levels are high. We found a similar fluctuation in the mechanical pain sensitivity across the estrous cycle in mice; the pain susceptibility was lower during diestrus phase compared to that during the estrus phase of the cycle. Other studies have found similar fluctuations in the susceptibility to thermal stimuli and visceromotor pain during estrous cycle (Frye et al., 1992; Martínez-Gómez et al., 1994; Ji et al., 2008).

Several prior studies have evaluated pain modulating effects of progesterone, but no study had focused on the effects dependent on PR activation. Many of these studies reported a pain-suppressing action of progesterone (Coronel et al., 2011; Verdi et al., 2013; Coronel et al., 2014; Coronel et al., 2016b; Coronel et al., 2017; Hornung et al., 2020); however, other studies found that progesterone treatment was ineffective in suppressing neuropathic pain (Dableh and Henry, 2011; Ungard et al., 2020). The pain-promoting action of PR activation uncovered here may oppose the analgesic effects of allopregnanolone which could make progesterone treatment ineffective in suppressing pain. Furthermore, Ungrad et al found that progesterone was robustly protective in males with neuropathic pain, but had only a moderate effect in females. We found that PR agonist segesterone did not exert a pain-promoting effect in males and pre-pubertal females, raising the possibility that expression of PRs and down-stream signaling molecules could be distinct in adult females and in adult males and juvenile females.

PR deletion blocked the pain-promoting effects of progesterone and segesterone. Two prior studies have found that PR antagonists RU-486 and ICI 182,780 could protect from hypersensitivity associated with neuropathic pain. In one study, RU-486 treatment blocked mechanical allodynia and thermal hyperalgesia induced by spinal nerve injury (Takasaki et al., 2005). Although, this study correlated the improvement in pain to glucocorticoid receptors (GRs), RU-486 also blocks PR actions. In another study ICI 182,780 an anti-PR and anti-estrogen receptor antagonist prevented neuropathic pain (Kondo et al., 2006). However, in both these studies the protective effects were noted soon after the antagonist administration, raising the possibility that distinct mechanisms may regulate these acute pain-alleviating effects rather than the slow-emerging effects seen here.

Besides regulating the pain threshold in healthy animals, we have also uncovered a potential role of PRs in regulating migraine-accompanying sensory hypersensitivity. The perimenstrual period is associated with a heightened susceptibility to migraine headaches, whereas some women experience headaches exclusively during menstruation (MacGregor and Hackshaw, 2004; Lay and Payne, 2007; Vetvik et al., 2013). The role of progesterone in regulating perimenstrual migraine is underexplored. In one clinical trial performed in a limited number of patients, exogenous progesterone administration did not suppress migraine headaches (Somerville, 1971; Anne MacGregor, 2004). Based on the findings presented here, we propose that the luteal rise in progesterone would activate PRs. However, the pain-promoting effect of this activation would remain suppressed until the strength of inhibitory neurotransmission is maintained by high levels of circulating progesterone. With the perimenstrual decline in progesterone levels, the inhibitory neurotransmission would weaken and the effects of PR activation would emerge. A single injection of progesterone caused an acute elevation in circulating progesterone levels which returned to the baseline levels after 16 hours. However, this single administration of progesterone was sufficient to lower the pain threshold for the duration of testing (4 days). Thus, the PR-regulated effects could last even after the progesterone levels had declined. We used paw withdrawal threshold as a measure of pain here. Other studies have used periorbital pain threshold as a measure of migraine-accompanying pain (De Logu et al., 2019; Mason et al., 2021; Bertels et al., 2022), which could be a better correlate of migraine-accomapnying pain in humans. However, anti-migraine therapies also block the peripheral allodynia induced by NTG (Pradhan et al., 2014; Tipton et al., 2015; Moye et al., 2021).

Progesterone receptor gene polymorphism (PROGIN), which is predicted to affect the receptor function is seen in migraine patients. However, whether these mutations exacerbate or alleviate this condition is unclear (Colson et al., 2005; Corominas et al., 2009; Joshi et al., 2010; Schürks et al., 2010; Rodriguez-Acevedo et al., 2013; Palmirotta et al., 2015). The diverse study population and distinctions in clinical characterization of patients could have influenced these results. The current study in experimental animals with controlled hormone levels and treatment of a specific PR agonist found that PR activation could exacerbate this disorder.

Headaches are a common side effect of the oral contraceptives in women, and the use of combined hormonal contraceptives may increase the risk of migraines (Kudrow, 1975; Massiou and MacGregor, 2000; Aegidius et al., 2006; Teepker et al., 2011). On the other hand, retrospective studies have found reduction in the number of headache and migraine days in women taking progestin-only contraceptive pill (Nappi et al., 2011; Merki-Feld et al., 2013; Morotti et al., 2014). Although, this finding seems at odds with the role of PRs uncovered here, estrogens induce PR expression, whereas progesterone exerts a suppressive effect (Guerra-Araiza et al., 2003; Mani and Portillo, 2010). A potential reduction in the PR expression together with allopregnanolone-mediated analgesic effects in these women may improve headaches. In one study average pain tolerance was more in healthy women taking combined oral contraceptives compared to that in men or menstruating women (Reddy B, 2022). This indicates that hormonal contraceptives may protect healthy women from pain. However, whether the contraceptives have a similar effect in conditions like migraine or chronic neuropathic pain in which activity of the pain matrix is altered remains unexplored.

These studies were informed by the emerging studies elucidating a role of PRs in regulating neuronal excitability and glutamatergic neurotransmission. We have found that PR activation potentiates glutamatergic transmission through AMPA receptors and neuronal excitability (Joshi et al., 2018; Shiono et al., 2021). The pain matrix is composed of sensory neurons of the dorsal horn, periaqueductal grey, thalamus, somatosensory, insular, cingulate, and prefrontal cortices, and amygdala (May, 2009). Additionally, trigeminal ganglia, spinal trigeminal caudal nucleus, and hypothalamus are also associated with the migraine pain (Goadsby et al., 2017). A potential PR-regulated the increase in excitability and/or strengthening of glutamatergic transmission of one or more components of the pain matrix could increase pain sensitivity. This regulation could happen directly or indirectly. PRs are expressed in the cortices, amygdala, and hypothalamus (Maclusky and McEwen, 1978; Parsons et al., 1982; Cerbon et al., 1989; Hagihara et al., 1992; Guerra-Araiza et al., 2000; Shiono et al., 2021), and may also be present in the spinal trigeminal caudal nucleus (TNC) (Haywood et al., 1999). Thus in depth characterization of PR expression in the pain matrix and understanding the effect of PR activation on the activity of regions could help identify nodes that are potential targets of PRs. Furthermore, we determined the paw withdrawal threshold as a measure of pain sensitivity in these studies. Follow-up studies evaluating cephalic allodynia in terms of periorbital pain threshold and light sensitivity will be critical in distinguishing the role of PRs in regulating the activity of trigeminal pain pathway and/or spinothalamic pain pathways that regulate cephalic and extracephalic nociception. Additional mechanisms that may regulate pain downstream of the progesterone-PR signaling may involve anti- and pro-inflammatory molecules. Two prior studies have found that the anti-inflammatory actions of progesterone are blocked in the PRKO mice (Coronel et al., 2011; Labombarda et al., 2015). PRs also interact with synaptic proteins in the hypothalamus (Acharya et al., 2017), and similar actions at other synapses in the pain pathway may also be important.

In conclusion, we have uncovered a novel pain-promoting effect of progesterone that is slow to emerge. This effect depends on PR expression. PRs also seem to regulate migraine-accompanying sensory hypersensitivity. Thus, PRs could represent a potential target for the treatment of migraine and other chronic pain conditions that disproportionately affect women of reproductive age.

## Acknowledgements

This study was supported by the National Institutes of Health (NIH) grants R01 NS 110863 to SJ and R01NS120945, R37NS119012 to JK. The content is solely the responsibility of the authors and does not necessarily represent the official views of the National Institutes of Health. We thank Dr. Amynah Pradhan, Washington University in St. Luis for critical comments and Elizaveta Mangutov, Washington University in St. Louis for guidance on the mechanical pain threshold measurements.

## Data availability

The datasets generated and analyzed in this work are available from the corresponding author upon reasonable request.

## Supplementary material

Supplementary figure 1: NTG did not cause a further reduce pain threshold in animals that were pretreated with a high dose segesterone (10 mg/kg). The values represent mean and SD, n= 5 each. **(A)** The pain threshold in the animals that were pretreated with vehicle (cyclodextrin) in the place of segesterone, black bars represent animals treated with vehicle for NTG (a solution of 30% alcohol and 30% propylene glycol, diluted 1:1 with saline) and red bars represent pain threshold in the animals that received NTG, p=0.003 ordinary one-way ANOVA. The values of post-hoc Holm-Šídák’s multiple comparisons test are marked on the graph. **(B)** The pain threshold in segesterone-pretreated mice after administration of vehile or NTG, p= 0.029 ordinary one-way ANOVA.

